# Prion protein quantification in cerebrospinal fluid as a tool for prion disease drug development

**DOI:** 10.1101/295063

**Authors:** Sonia M Vallabh, Chloe K Nobuhara, Franc Llorens, Inga Zerr, Piero Parchi, Sabina Capellari, Eric Kuhn, Jacob Klickstein, Jiri Safar, Flavia Nery, Kathryn Swoboda, Stuart L Schreiber, Michael D Geschwind, Henrik Zetterberg, Steven E Arnold, Eric Vallabh Minikel

## Abstract

Reduction of native prion protein (PrP) levels in the brain is an attractive and genetically validated strategy for the treatment or prevention of human prion diseases. However, clinical development of any PrP-reducing therapeutic will require an appropriate pharmacodynamic biomarker: a practical and robust method for quantifying PrP, and reliably demonstrating its reduction, in the central nervous system (CNS) of a living patient. Here we evaluate the potential of enzyme-linked immunosorbent assay (ELISA)-based quantification of human PrP in human cerebrospinal fluid (CSF) to serve as a biomarker for PrP-reducing therapeutics. We show that CSF PrP is highly sensitive to plastic adsorption during handling and storage, but its loss can be minimized by addition of detergent. We find that blood contamination does not affect CSF PrP levels, and that CSF PrP and hemoglobin are uncorrelated, together suggesting that CSF PrP is CNS-derived, supporting its relevance for monitoring the tissue of interest and in keeping with high PrP abundance in brain relative to blood. In a cohort with controlled sample handling, CSF PrP exhibits good within-subject test-retest reliability (mean coefficient of variation 13% in samples collected 8-11 weeks apart), a sufficiently stable baseline to allow therapeutically meaningful reductions in brain PrP to be readily detected in CSF. Together, these findings supply a method for monitoring the effect of a PrP-reducing drug in the CNS, enabling the development of prion disease therapeutics with this mechanism of action.

## Introduction

Prion disease — a fatal and incurable neurodegenerative disease — is caused by misfolding of the prion protein (PrP), encoded by the gene *PRNP*^1^. PrP is a well-validated drug target for prion disease: knockout animals are invulnerable to prion infection^2^, heterozygous knockouts have delayed onset of disease^3^, and post-natal depletion of PrP can delay or prevent prion disease^4,5^. Total knockout is tolerated in mice^6,7^, cows^8^, and goats^9,10^, and healthy humans with one loss-of-function allele of *PRNP* have been identified^11^. Therefore, candidate therapies for prion disease may seek to lower PrP levels in the brain. Similar approaches are being explored in other neurodegenerative diseases, with promising preliminary results in humans^12,13^.

Clinical trials of PrP-lowering therapies will be enhanced by early determination of whether PrP is indeed being lowered effectively at a tolerated dose. The brain is the target tissue for any prion disease therapeutic, but is difficult to monitor directly. Cerebrospinal fluid (CSF) is produced by the choroid plexus of the ventricles, flows in and around the spinal cord and is in intimate contact with interstitial fluid of brain parenchyma. CSF more closely reflects the biochemistry of the brain than blood or any other accessible tissue, and is obtainable through a minimally invasive lumbar puncture (LP). PrP levels in CSF range from tens to hundreds of ng/mL, within the range of standard protein detection assays. Multiple groups have reported successful detection of PrP in human CSF using ELISA assays, including the one currently commercially available human PrP ELISA kit, the BetaPrion^®^ ELISA assay^14–18^ (Analytik Jena, Leipzig, Germany). The assay is best described as measuring total PrP, which is the variable of interest for PrP-lowering therapeutics (see Discussion).

Informed by FDA’s 2013 Draft Guidance on Bioanalytical Method Validation^19^ we assessed the technical performance of the BetaPrion^®^ ELISA assay across *N*=225 human CSF samples spanning a range of diagnoses. We then used this assay to investigate the biological suitability of CSF PrP as a pharmacodynamic biomarker for PrP-reducing therapeutics.

## Results

### The BetaPrion^®^ Human PrP ELISA quantifies total CSF PrP reproducibly, precisely, sensitively, and selectively

We assessed the assay’s precision, sensitivity, selectivity and reproducibility by analyzing *N*=225 human CSF samples from symptomatic prion disease patients, pre-symptomatic prion disease mutation carriers, non-prion dementia patients, and normal pressure hydrocephalus (NPH) patients as well as other non-prion controls (Table S1) across 41 plates. The results broadly support the technical suitability of this assay for reliable quantification of CSF PrP (Table 1 and Figure S1).

**Table 1.** The technical performance of the BetaPrion^®^ human PrP ELISA assay supports reliable quantification of PrP in human CSF. Abbreviations: coefficient of variation (CV); lower limit of quantification (LLOQ); amino acid analysis (AAA).

| Experiment | Results |
| --- | --- |
| Within-plate technical replicate reproducibility (same dilution) | CV = 8% |
| Within-plate technical replicate reproducibility (all dilutions) | CV = 11% |
| Between-plate technical replicate reproducibility | CV = 22% in an interplate control sample run on 17 plates on different days (see Supplementary Discussion). |
| Sensitivity | LLOQ is 3-5× the blank signal, depending on the platereader used. |
| Selectivity | Non-reactive for recombinant mouse PrP, rat CSF and cynomolgus monkey CSF (consistent with one amino acid mismatch in the reported detection antibody epitope<sup>17</sup>), artificial CSF and protease-digested CSF. |
| Dilution linearity | Linear across two samples and five dilutions. See Figure S1A. |
| Spike recovery | Using AAA-quantified recombinant HuPrP23-230 as a standard, spike recovery of recombinant PrP in CSF was 90% across five concentrations. Titration of a high PrP CSF sample into a low PrP sample resulted in linear recovery. See Figure S3. |
| Standard curve reproducibility | CV < 10% at all six non-zero standard curve points, across five replicates. See Figure S1. |

In assessing within-plate variability we discerned plate position effects for control samples, with a mild but significant downward trend from upper left to lower right (Figure S2). Comparison of the kit standard curve to a standard curve made from recombinant human prion protein quantified by amino acid analysis (AAA) yielded systematic differences, with implications for kit use for absolute versus relative quantification of PrP (Figure S3B; see Discussion).

### Standardized storage and handling are essential to reliable quantification of CSF PrP

PrP was measurable by ELISA in all *N*=225 CSF samples analyzed, including in CSF from individuals with 13 different genetic prion disease mutations (Figure S4A-B, Table S1). Across all CSF samples analyzed, PrP levels varied by over two orders of magnitude (Figure S4A), ranging from 1.9 to 594 ng/mL. PrP was reduced in individuals with symptomatic prion disease, as previously reported^14,15,17,18,20^. Within matched cohorts containing individuals with prion disease, however, diagnostic category (non-prion, presymptomatic genetic, symptomatic genetic, and sporadic prion disease) explained only a minority of variance in CSF PrP level (adjusted R^2 = 0.23, *P* < 1 × 10^−7^, linear regression). After excluding individuals with symptomatic prion disease, PrP still differed significantly between the various cohorts included in our study, and within-cohort variation was also dramatic (Figure S4C; mean ~20-fold difference between highest and lowest sample within a cohort). These observations led us to search for other factors that might contribute either to biological or pre-analytical variability. CSF PrP was correlated with age (Figure S4D), but among our samples age is confounded with cohort, diagnosis, and likely many unobserved variables, making it unclear whether this correlation is biologically meaningful. CSF PrP did not differ according to sex (Figure S4E), and exhibited no lumbar-thoracic gradient over serial tubes collected from the same LP (Figure S4F-G). After noticing that PrP levels appeared lower in smaller aliquots of the same CSF sample (Figure S5A), we hypothesized that differences in sample handling might be one major source of variability in observed CSF PrP levels.

It is known that other neurodegenerative disease-associated amyloidogenic proteins have a high affinity for plastics^21–23^, but PrP’s stability under different handling conditions has not previously been systematically investigated. To assess PrP’s susceptibility to differential CSF sample handling, we subjected aliquots of a single CSF sample to variations in 1) number of transfers between polypropylene storage tubes, 2) amount of exposure to polypropylene pipette tips, 3) storage aliquot size, 4) storage temperature, and 5) number of freeze-thaw cycles (Figure 1A). Strikingly, increased plastic exposure in the first three conditions dramatically reduced measurable PrP in solution (Figure 1A). To promote PrP solubility in our samples, we experimented with adding small amounts of 3-[(3-Cholamidopropyl)dimethylammonio]-1-propanesulfonate hydrate (CHAPS), a common zwitterionic surfactant known to enhance protein solubility in multiple contexts^24–26^. Addition of 0.03% CHAPS prior to aliquotting minimized PrP loss to plastic across most manipulations (Figure 2A). For instance, transferring a CSF sample to a new microcentrifuge tube three times eliminated at least 73% of detectable PrP (*P* < 1 × 10^−6^, two-sided t test) without CHAPS, but only 7.1% (*P* = 0.37) of PrP was lost in the presence of 0.03% CHAPS. Addition of CHAPS also increased total PrP recovery, presumably by preventing loss to the single polypropylene tube and tips used for plating samples (Figure S5), and was effective against loss to multiple plastics but not glass (Figure 1C). Storing CSF at room temperature for 24 hours or subjecting samples to three freeze-thaw cycles had a less dramatic impact on PrP that did not appear to be affected by CHAPS (Figure 1A-B and Figure S5D-E).

**Figure 1.**
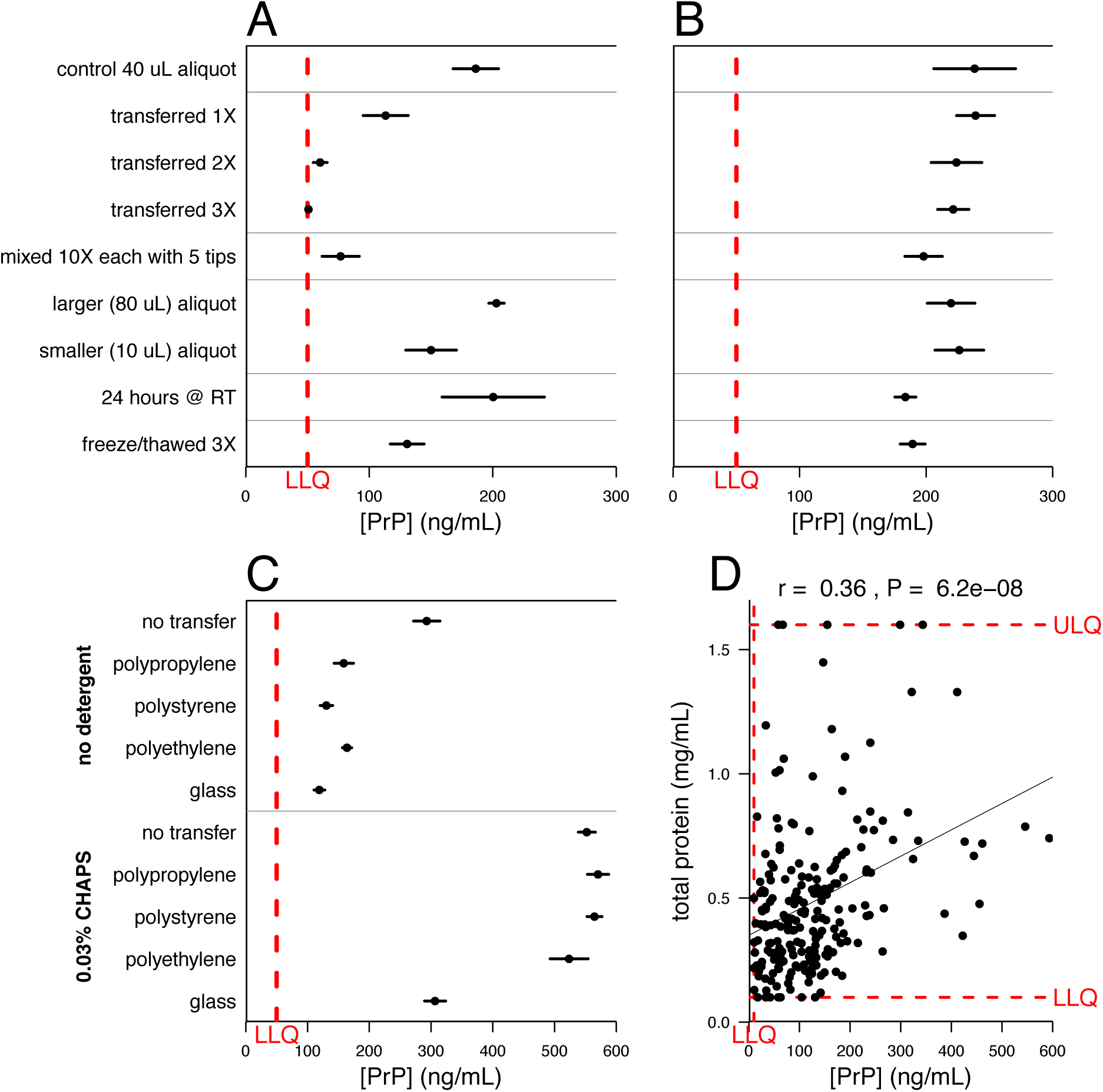
Storage and handling can dramatically reduce the amount of PrP detected in CSF samples unless appropriate measures are taken. In A-C, dots represent mean and line segments represent 95% confidence intervals across 4 to 7 aliquots of the same sample, each measured in duplicate at a 1:50 dilution. In D, dots represent mean of measurements within dynamic range, among 2 dilutions with 2 technical replicates each. A. Increased polypropylene exposure substantially reduces detectable PrP. B. Addition of 0.03% CHAPS detergent to samples increases PrP recovery and consistently mitigates PrP loss to plastic. C. Addition of CHAPS (bottom) increases total PrP recovery and shows similar rescue across plastics, but substantial PrP loss is still observed upon storage in glass. D. Across 217 CSF samples, total protein levels and PrP levels were modestly correlated (Spearman’s rank correlation coefficient = 0.36, P=6.2×10^−8^). In A-C, dots represent mean and line segments represent 95% confidence intervals across 4 to 7 aliquots of the same sample, each measured in duplicate at a 1:50 dilution. In D, dots represent mean of measurements within dynamic range, among 2 dilutions with 2 technical replicates each.

**Figure 2.**
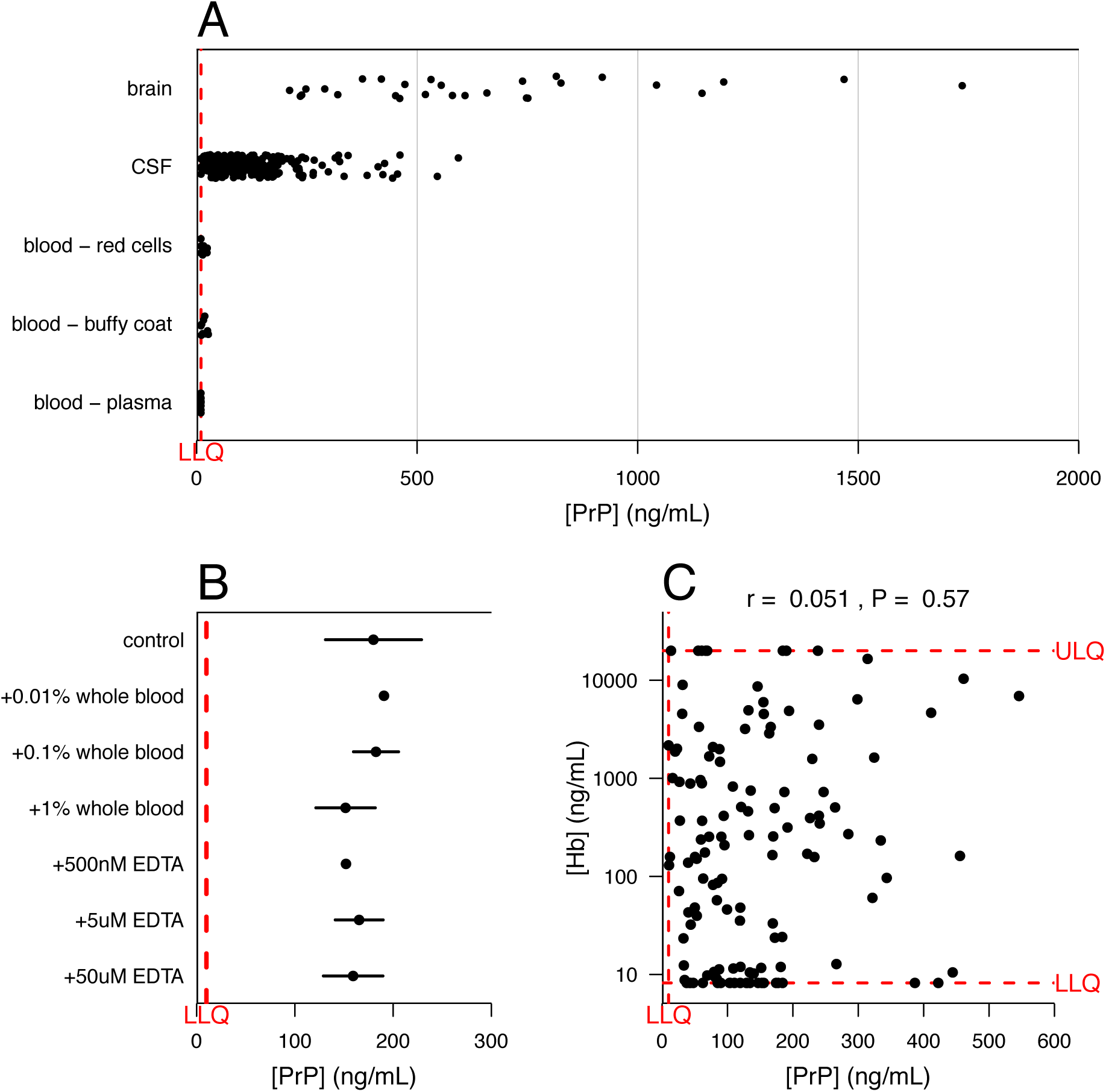
Blood PrP contributes negligibly to the PrP detected in CSF. A. PrP levels were compared by ELISA in N=28 postmortem human brain samples, three blood fractions from N=8 individuals each, and all N=225 CSF samples analyzed in the present study. PrP is abundant in a range of human brain regions, undetectable in human plasma, and is detectable in the red cell and buffy coat fractions only at low levels compared to PrP in CSF. B. Spiking whole blood into CSF up to 1% by volume does not impact measured PrP. C. Across N=128 CSF samples spanning multiple cohorts and diagnostic categories, hemoglobin and PrP levels in CSF are uncorrelated. In A and C, dots represent mean of measurements within dynamic range, among 2 technical replicates per dilution. In A-C, dots represent mean and line segments represent 95% confidence intervals across 2 to 3 aliquots of the same sample.

We also investigated the relationship between measured PrP and total protein in *N*=217 samples, using the DC total protein assay. Across all samples analyzed, a modest correlation (r = 0.36, Spearman rank test, *P* < 1 × 10^−7^) between PrP and total protein was observed (Figure 1D), which may reflect either a biological phenomenon, or simply the ability of higher ambient protein levels to serve a blocking function that partially offsets PrP loss by adsorption. In support of the latter interpretation, addition to CSF of 1 mg/mL bovine serum albumin increased recovery of PrP (Figure S5F), though it was less effective than CHAPS at preventing loss due to transfers.

### PrP in CSF is CNS-derived and unlikely to be confounded by blood contamination

CSF PrP is an informative tool in prion disease only insofar as it is a faithful proxy for PrP levels in the CNS, the relevant target for any future therapeutic. CSF proteins derive from two major sources, CNS and blood, with proportional contribution driven by relative tissue abundance of a given protein^27,28^. Blood proteins may enter CSF either through passive diffusion as CSF flows along the spinal cord^29^, or artifactually if blood from a traumatic lumbar puncture contaminates the collected CSF. To assess the contribution of blood-derived PrP to overall CSF PrP, we compared PrP levels across brain samples and red blood cell, buffy coat and plasma fractions of blood from non-neurodegenerative disease control individuals, versus all of the CSF samples in our study (Figure 2A). Among blood fractions, PrP was most consistently detected in buffy coat, in keeping with reports that blood PrP emanates chiefly from platelets^30,31^; we also detected PrP above the lower limit of quantification in some red cell samples, but never in plasma. As the average PrP concentration in all three blood fractions was still well below that in brain and was lower than that in 96% of CSF samples analyzed, the risk of confounding signal from blood-derived PrP appears negligible. Consistent with this conclusion, spiking whole blood into CSF at up to 1% (v/v) did not increase the detected PrP (Figure 2B). Finally, as a proxy for blood contamination we measured hemoglobin levels in *N*=128 CSF samples and observed no correlation between CSF hemoglobin and CSF PrP (Figure 2C). Variation in hemoglobin levels also failed to confound the test-retest reliability of CSF PrP (Figure S6). From these lines of evidence we conclude that the PrP detected in CSF is overwhelmingly derived from the CNS.

### CSF PrP levels in individuals are stable on short-term test-retest

In order for CSF PrP levels to serve as a meaningful biomarker, they must be stable enough in one individual over time that a drug-dependent reduction could be reliably detected. We quantified PrP in pairs of CSF samples collected from nine individuals — placebo-treated controls with non-prion dementia — who had undergone two fasting morning lumbar punctures at 8-11 week intervals in the context of a clinical trial^32^. LPs were performed according to a standardized protocol by a single investigator, and samples were subsequently processed uniformly. Under these highly controlled conditions, the mean CV between timepoints for a given participant was reasonably low at 13% (Figure 3). Higher CVs of 33% - 41% were observed in three other cohorts where sample handling appears to have been less uniform (Supplementary Discussion and Figure S7), consistent with PrP’s susceptibility to pre-analytical perturbations (Figure 1).

**Figure 3.**
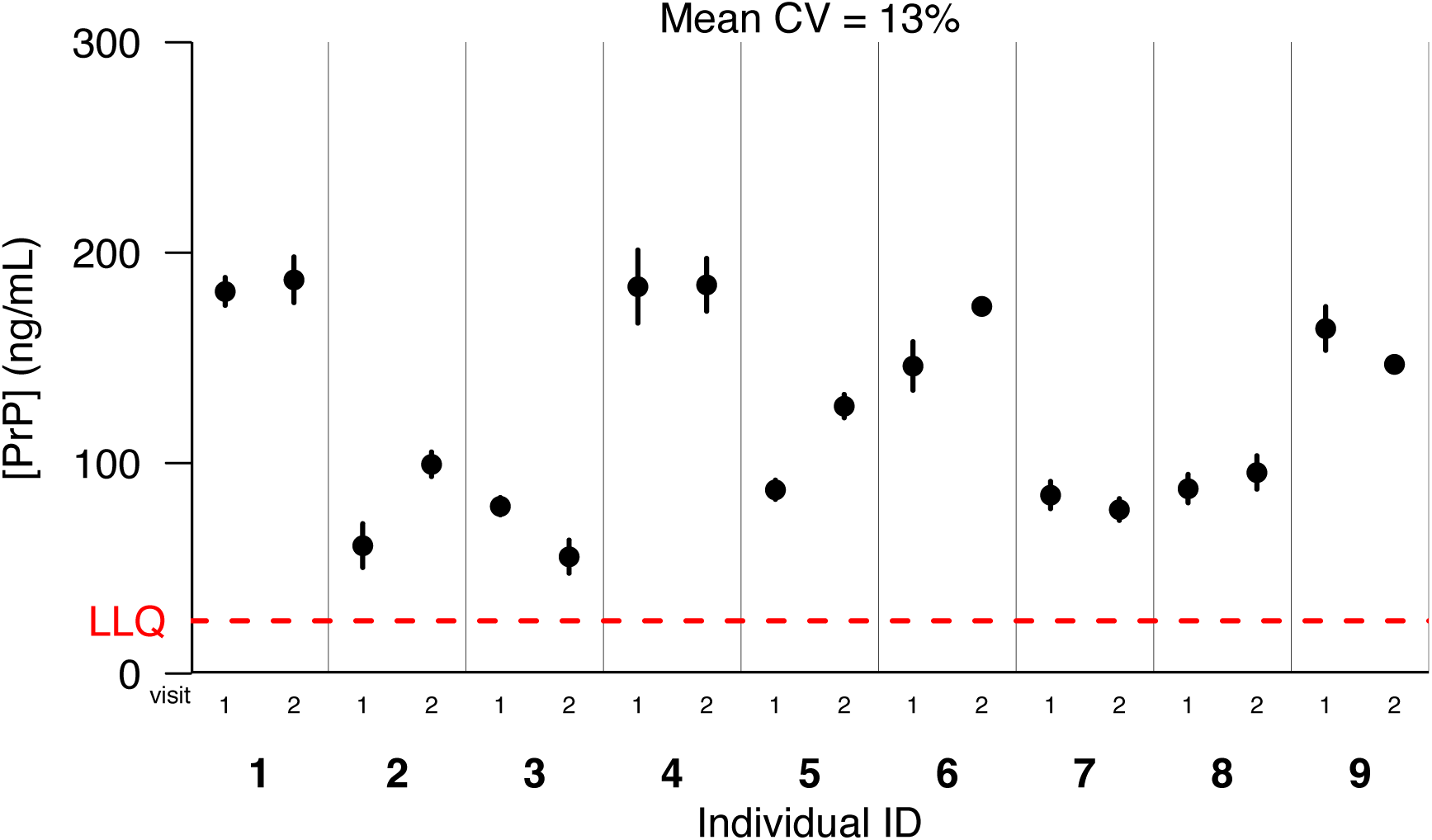
Test-retest stability of CSF PrP. Uniformly processed CSF samples were provided from a past clinical trial, from placebo-treated individuals with mild, non-prion cognitive impairment. Fasting morning lumbar punctures were performed by one investigator on nine individuals then repeated at an interval of 8-11 weeks. Dots represent means, and line segments 95% confidence intervals, of measurements within dynamic range among 2 dilutions with 2 technical replicates each.

## Discussion

Here we present evidence supporting CSF PrP quantification as a tool for clinical trials of PrP-lowering therapeutics. We establish that CSF PrP is sensitive to multiple factors that may be encountered during handling and processing, and that the addition of 0.03% CHAPS detergent mitigates the most dramatic such factor by minimizing PrP loss to plastic. With use of appropriate protocols, CSF PrP can be measured reproducibly and with favorable test-retest reliability, with a mean CV of 13% over 8-11 weeks in one cohort. CSF PrP is CNS-derived, rather than blood-derived, suggesting it should change in response to lowering of PrP in the brain.

The above attributes suggest that CSF PrP will be a useful pharmacodynamic biomarker in the development of PrP-lowering therapeutics. However, our findings regarding PrP’s sensitivity to handling and processing factors demonstrate that an optimized protocol for CSF collection and processing will need to be closely followed for samples to be meaningfully compared. To this end, the protocol we are now using to collect such samples is detailed in Figure S8. For maximum protection of PrP from plastic adsorption, we propose addition of 0.03% CHAPS immediately upon transfer of CSF from the initial lumbar puncture syringe, prior to aliquotting or freezing. Detergent type and level were chosen for compatibility with our downstream ELISA and mass spectrometry assays. As shown in Figure S8, we are reserving aliquots of CSF without additive for future use in detergent-incompatible assays, but do not recommend use of such aliquots for PrP quantification. As CHAPS did not offer complete protection from plastic adsorption, and may not affect temperature-related insults, even in the presence of detergent these pre-analytical variables should still be a) minimized, b) closely tracked for all samples, and c) standardized for samples across which PrP levels will be compared.

In addition to best practices for sample handling, our experiments suggest best practices for assay use. In light of subtle plate position effects (Figure S2), samples intended for comparison, such as serial samples from one subject, should be co-located on the ELISA plate, and/or plate position should be adjusted for using standard curves or control samples. Our comparison of the kit standard curve to a standard curve made from recombinant human prion protein quantified by amino acid analysis (AAA) suggests that the kit may be most useful for relative quantification of PrP (for example, before and after administration of a PrP-lowering treatment) rather than absolute quantification (Figure S3B).

As previously mentioned, PrP levels in CSF as measured by ELISA have been reported to be reduced on the order of half in symptomatic prion disease patients^15,17,18^, and this phenomenon is reflected in our samples as well (Figure S4). Multiple plausible biological mechanisms could explain these findings: incorporation of PrP into insoluble plaques^33,34^, internalization of misfolded PrP in the endosomal-lysosomal pathway^35^, and post-translational downregulation of PrP as a function of disease^36^. It is therefore possible that an intrinsic reduction in CSF PrP in the course of symptomatic disease could confound the use of PrP as a biomarker for the activity of PrP-lowering drug tested in a symptomatic population. Although it is important to be aware of this potential limitation, symptomatic patients are not the population most in need of such a biomarker. The signature rapid clinical decline associated with active disease has enabled several previous clinical trials to be conducted in symptomatic cohorts based on cognitive or survival endpoints^37–43^, and future trials may be further benefit from the use of real-time quaking induced conversion (RT-QuIC) to detect misfolded prion “seeds” in symptomatic patient CSF^44–46^. Instead, the population best positioned to benefit from a CSF PrP pharmacodynamic biomarker in conjunction with a PrP-lowering drug is the population of presymptomatic individuals carrying high-penetrance genetic prion disease mutations. Our experiments confirm that PrP is measurable in carriers across a variety of mutations. They also support the hypothesis that target engagement and achievement of a meaningful proximal biological effect by a PrP-lowering drug candidate could be observed through quantification of PrP in CSF from serial lumbar punctures in such individuals, a hypothesis that will need to be tested in a clinical trial once such a drug candidate is available.

Our study has several limitations. First, ELISA relies upon two epitopes being present and properly folded, and is thus vulnerable to confounding from misfolding or native proteolytic events. We are presently working to develop a targeted mass spectrometry-based orthogonal method for CSF total PrP quantification. Second, although we have established that CSF PrP is quantifiable in genetic prion disease patients and has good test-retest reliability in a cohort of patients with non-prion dementia, when we embarked on the present study we did not have access to short-term test-retest samples from presymptomatic genetic prion disease mutation carriers. To address this shortfall, in summer 2017 we launched a clinical research study at Massachusetts General Hospital to recruit presymptomatic individuals with *PRNP* mutations, and controls, for two lumbar punctures at an 8- to 16-week interval^47^. This study is following the collection and processing protocol specified in Figure S8. We hypothesize that with this protocol in place, test-retest reliability in this population will prove sufficient to enable future clinical trials monitoring CSF PrP before and after administration of a PrP-lowering drug. Third, the samples analyzed here were re-used after collection for other research or clinical purposes, meaning that in most cases we cannot fully account for how the samples were handled prior to our receipt of them. Thus, our numbers may exaggerate the inter-individual variation in CSF PrP in the population. The question of whether the large observed inter-individual variability in PrP CSF levels indicates true biological variability or handling artifacts will be addressed by the uniformly processed samples currently being collected through our clinical study.

In recent studies of potential Alzheimer’s disease and Huntington’s disease biomarkers, the goal has been detection of pathological molecules such as Aβ oligomers or mutant huntingtin protein that are thought to be causative or otherwise indicative of their respective disease processes^22,48^. Our goal differs in that native PrP is present in all humans, and in its native state is not pathogenic; it is present and measurable in healthy individuals. PrP is an attractive drug target in prion disease because it is positioned upstream on the disease pathway, and as the substrate for conversion to pathogenic misfolded prions, presents a shared target among all prion disease subtypes. Further, unlike the misfolded prions that derive from it, PrP is structurally well-characterized and can be targeted genetically; approaches to protectively reducing its levels could intervene at the level of DNA, RNA or protein. Our findings should enable clinical development towards the realization of this well-supported therapeutic strategy.

Moreover, this biomarker may help to empower a heretofore-unexplored route for drug development and trials in healthy, at-risk individuals. The increasing availability of large genetic datasets is enabling improved estimation of the penetrance associated with disease-associated *PRNP* mutations^11^. Individuals facing 90% or greater lifetime risk of genetic prion disease can be reliably identified years or decades in advance of onset. Such carriers lack any overt phenotype, and to date no reliable change indicative of prodromal disease has been reported in this population by imaging or biochemical analysis. In addition, even if such a marker were to be found, it would be useful only once the prodromal disease process were already underway, when the greatest opportunity for meaningful intervention in at-risk individuals may have already passed. Preclinical evidence strongly indicates that regardless of mechanism of action, the potency of anti-prion therapeutics scales with time of intervention relative to disease course, with prophylactic administration prior to any molecular pathology offering greater benefit in delaying disease^49–52^. In the context of prevention trials in healthy carriers, it is possible that CSF PrP will be critical not just as a marker of target engagement, but as a surrogate endpoint. Because following pre-symptomatic individuals to a clinical endpoint appears infeasible^53^, lowering CSF PrP has been proposed as a surrogate endpoint meriting Accelerated Approval^54^. Continued study of CSF PrP will be critical to steering future treatment trials towards a preventative paradigm and to honoring the precious opportunity for preemptive intervention provided by predictive genetic testing.

## Methods

### Cerebrospinal fluid samples

De-identified human CSF samples were provided by multiple clinical collaborators and included both unpublished and previously published cohorts^32,55^. Samples were shipped on dry ice and stored at −80°C. Prior to use, samples were thawed on ice and centrifuged at 2,000 × g (at 4°C). Ninety percent of the volume was pipetted into a new tube to separate supernatant from cellular or other debris, aliquotted into new polypropylene storage tubes and refrozen at −80°C. For indicated samples, 0.03% CHAPS detergent by volume (final concentration, from a 3% CHAPS stock) was pre-loaded into the supernatant receiving tube prior to the post-centrifugation transfer, then mixed into the sample by gentle pipetting prior to aliquotting.

### Quantification of human PrP in CSF, brain tissue and blood using the BetaPrion^®^ human PrP ELISA kit

Across experiments, PrP was quantified using the BetaPrion^®^ human PrP ELISA kit (Analytik Jena, cat no. 847-0104000104) according to the manufacturer’s instructions. This sandwich ELISA is configured in 96-well format and relies on an apparently conformational human PrP (HuPrP) capture antibody and a horseradish peroxidase (HRP)-conjugated primary detection antibody to HuPrP residues 151-180^17^. In brief, samples were diluted into blocking buffer (5% BSA and 0.05% Tween-20 in PBS, filtered prior to use) at concentrations ranging from 1:100 to neat depending on the anticipated PrP content of the sample type. All samples were plated in duplicate. Lyophilized standards and kit reagents were diluted fresh for same-day use, with the exception of wash buffer and blocking buffer, excess of which were stored at 4°C for reuse within 4 weeks. The assay format is 96-well comprised of twelve modular 8-well strips which enabled partial plates to be run in some cases. Following all add and incubation steps the absorption per well was read in either a SpectraMax or FluoStar Optima plate reader at 450 nm with 620 nm absorbance also monitored as baseline. Data was exported as a text file and analyzed in R.

Unknown CSF samples were run at two dilutions each (typically 1:10 and 1:50). Only one out of 225 CSF samples analyzed fell below the range of the assay’s lower limit of detection (1 ng/mL final) at a 1:10 dilution, and was re-run neat, yielding a result of 1.9 ng/mL. Except where noted, samples were run in technical duplicate at two dilutions, and error bars represent 95% confidence intervals around the mean.

Human brain samples were obtained from the Massachusetts Alzheimer’s Disease Research Center (ADRC; *N*=26 samples from 5 different control individuals without neurodegenerative disease, with post-mortem intervals of 23-72 hours, representing diverse cortical and subcortical regions) and from the National Prion Disease Pathology Surveillance Center (*N*=2 samples of frontal cortex from non-prion controls) homogenized in PBS with 0.03% CHAPS at 10% weight/vol in 7mL tubes (Precellys no. KT039611307.7) using a MiniLys tissue homogenizer (Bertin no. EQ06404-200-RD000.0) for 3 cycles of 40 seconds at maximum speed. The resulting 10% brain homogenates were diluted 1:10 and 1:100 in blocking buffer for ELISA.

Human blood fractions were obtained from Zen-Bio (3 fractions – red blood cell, buffy coat, and plasma – from 8 individuals each), 0.03% CHAPS was added, and samples were then mixed either by pipetting up and down or by homogenization in a MiniLys using the same protocol described above. Blood fractions were diluted 1:10 in blocking buffer for ELISA.

### Negative controls

Rat and cynomolgous monkey CSF (BioReclamation IVT; two samples each from two separate animals) and artificial CSF (Tocris no. 3525) were aliquotted and stored at −80°C. For protease-digested CSF, two CSF samples with 0.03% CHAPS (measured to contain 273 and 643 ng/mL PrP undigested) were digested with 5 µg/mL Proteinase K (WW Grainger Co. cat. no. 5000186667) at 37C for 1 hour, after which the digestion was halted with 4 mM PefaBloc (Sigma Aldrich cat. no. 11429868001) immediately prior to use in ELISA.

### Recombinant prion protein purification

For spike-in experiments and attempted detection of mouse recombinant PrP, in-house purified recombinant full-length human prion protein and mouse prion protein were purified from *E. coli* using established vectors (a generous gift from Byron Caughey’s laboratory at NIH Rocky Mountain Labs) according to established methods^56,57^. Protein concentration was determined by 280 nm absorbance on a NanoDrop, and by amino acid analysis (AAA) performed in duplicate (New England Peptide) after the addition of 0.03% CHAPS.

### Storage and handling experiments

For all storage and handling experiments, each condition was run in parallel on four identical aliquots made from one original CSF sample, and each aliquot was plated in duplicate. For all transfer experiments, 40 µL CSF aliquots were thawed on ice, then the full volume was transferred to a new 500 µL storage tube the indicated number of times and allowed to sit for a minimum of fifteen minutes in each tube. Where not otherwise indicated, tubes were polypropylene, and sample aliquots were 40 µL.

### Total protein assay

The DC total protein assay (Bio-Rad cat. no. 5000111) was used according to the manufacturer’s instructions to measure total protein across 217 CSF samples (all samples in this study except for the N=8 lumbar-thoracic gradient samples, Figure S4F-G). This assay, similar in principle to a Lowry assay, combines the protein with an alkaline copper tartrate solution and Folin reagent^58^. The protein reacts with copper in the alkaline medium, then reduces the Folin reagent to yield species with a characteristic blue color in proportion to abundance of key amino acids including tyrosine and tryptophan.

### Whole blood spike-in

Human whole blood (Zen-Bio) was spiked into parallel aliquots of a single CSF sample containing baseline mid-range PrP at 1%, 0.1%, or 0.01% per volume. EDTA spike-ins were performed in parallel to control for EDTA preservative carried in the blood sample. Samples were refrozen following spike-in then re-thawed for use to ensure lysis of cellular fractions prior to PrP quantification.

### Bethyl Laboratories Human Hemoglobin ELISA

Hemoglobin was quantified in 128 human CSF samples using the Human Hemoglobin ELISA kit (Bethyl Laboratories no. E88-134), according to the manufacturer’s instructions. Samples for this analysis spanned diagnostic categories including normal pressure hydrocephalus, non-prion dementia, symptomatic genetic and symptomatic sporadic prion disease. Samples were diluted 1:10 and 1:100 for most experiments, an in some cases 1:20 and 1:100. All samples were plated in duplicate.

### Blinding procedures

Assay operators were blinded to diagnosis for prion disease CSF cohorts. For test-retest cohorts, assay operators were blinded to test-retest pairing for Metformin trial samples and MIND Tissue Bank samples; pairing was known but collection order unknown for UCSF samples; pairing and order were known for Sapropterin trial samples.

### Statistics, data, and source code availability

All statistical analyses were conducted, and figures generated, using custom scripts in R 3.1.2. Raw data from platereaders, associated metadata, and source code sufficient to reproduce the analyses reported herein are publicly available at: https://github.com/ericminikel/csf_prp_quantification/

## Acknowledgments

This study was approved by the Broad Institute’s Office of Research Subjects Protection (ORSP-3587 and NHSR-4693). SV is supported by the National Science Foundation (GRFP 2015214731). EVM is supported by the National Institutes of Health (F31 AI122592). This work was supported by BroadIgnite and the Next Generation Fund at the Broad Institute of MIT and Harvard, Prion Alliance, anonymous donors, and an anonymous organization. Work in Germany was supported by Robert-Koch-Institute through funds of Federal Ministry of Health (grant no. 1369-341) to IZ and Spanish Ministry of Health-Instituto Carlos III (Miguel Servet-CP16/00041) to FL. UCSF sample collection was supported by NIH/NIA R01 AG-031189 and R01 AG-032289 and the National Center for Advancing Translational Sciences NIH UCSF-CTSI UL1 RR024131. We thank the Massachusetts Alzheimer’s Disease Research Center (ADRC, supported by NIH P50 AG005134) and the MassGeneral Institute for Neurodegenerative Disease (MIND) Tissue Bank for providing human brain and cerebrospinal fluid samples for this study. We thank Joanne Kotz for discussions. We are also grateful to the patients and their families who contributed samples to this research.

## Supplementary Discussion

### Technical parameters of the BetaPrion^®^ ELISA kit.

As noted in Table 1, for one sample included as an inter-plate control on 17 different plates, we observed an inter-plate CV of 22%. The 17 plates included in our analysis include plates from three different manufacturer lots, run by two different operators (SV and EVM), read on two different platereaders (Fluostar Optima and Spectramax), all of which factors may contribute to the variability we observed.

On an intra-plate basis, we also observed slightly higher variability when including dilutions than when only comparing replicates at a single dilution (CV=11% vs. 8%). Most samples were analyzed at two dilutions, 1:10 and 1:50, with two replicates each. In many cases, one dilution or the other fell outside the assay’s dynamic range, but among *N*=87 samples for which both the 1:10 and 1:50 dilutions had both replicates fall within the dynamic range of the assay (1 to 20 ng/mL final), the PrP level indicated by the 1:10 dilution was on average 3.5% higher than the 1:50 dilution.

### Plate position effects.

To assess whether plate position affects apparent PrP levels in ELISA, we ran two whole ELISA plates loaded with technical replicates of the same CSF sample (v1209 with 0.03% CHAPS). One plate was loaded with a single channel pipette taking 29 minutes (Figure S2A-B) and the other was loaded with a multichannel pipette taking 11 minutes (Figure S2C-D). A visually subtle, yet significant (*P* = 1.5e-14, linear regression), decline in apparent PrP level is seen across the plate. For instance, in Figure S2A, the ten replicates loaded last (wells G9-H6) are on average 22% lower than the ten replicates loaded first (wells A11-B8). Adjustment based on the standard curves abolishes this slope, and reduces the CV among technical replicates (Figure S2B and D).

### Spike recovery experiments.

While we ultimately achieved 90.5% recovery of recombinant human PrP spiked into CSF, this successful outcome was preceded by a number of experiments that usefully illuminate constraints of working with both the BetaPrion^®^ ELISA assay and CSF PrP as an analyte. In our first experiment, recombinant full-length human PrP with concentration orthogonally established by amino acid analysis (AAA) was spiked into two CSF samples previously established to have high and low baseline PrP. Compared to the expected recovery, the recombinant protein gave a much higher signal than expected, with 392-451%, over-recovery (Figure S3A). This surprising finding suggested to us that the concentration of PrP in kit standards may be lower in practice than the stated concentration. To test this hypothesis, we directly compared the kit standard curve to a matched standard curve prepared with our recombinant PrP. This experiment confirmed that kit standards appeared lower than AAA-quantified PrP standards by a factor of roughly 4 (Figure S3B). We conclude that kit standards, while technically reproducible, may most usefully inform relative rather than absolute quantification of PrP.

We next attempted to assess spike recovery in an internally consistent system by comparing recombinant PrP spiked into CSF to a recombinant PrP standard curve. We diluted recombinant PrP in CSF, then serially diluted into additional CSF to create a five-point series. The series of samples was re-frozen and measured by ELISA the next day. Under these intensive handling conditions, we observed only ~50% recovery even though the samples contained 0.03% CHAPS (Figure S3C). We hypothesized that the CHAPS additive, while helpful, could not fully protect against the high levels of plastic exposure involved in serial dilution of CSF. To test this hypothesis, we redid the experiment in C with special attention to protecting PrP from plastic adsorption. Recombinant PrP was diluted in blocking buffer to prepare a series of solutions at 100x the desired final concentrations of points in the spike series. These samples were then added to CSF aliquots at a 1:100 concentration, and used in a same-day ELISA experiment. With this level of attention to plastic exposure and the elimination of an additional freeze-thaw cycle relative to the standard curve, PrP was preserved near expected levels with 90.5% recovery observed (Figure S3D).

Finally, to assess recovery from a different angle, we titrated a high-PrP CSF sample into a low-PrP CSF sample at varying ratios, again ensuring minimal and consistent CSF handling. Under these conditions, we observed linear and proportional recovery of PrP (Figure S3E). These experiments provide additional evidence that the quality of PrP measurement afforded by the BetaPrion^®^ ELISA assay is dependent on appropriate sample processing.

### CSF aliquot size and PrP loss.

We observed that when working with experimental aliquots of CSF, lower volume aliquots appeared to have consistently lower PrP levels (Figure S5A). This effect is likely due to increased exposure of the sample to plastic due to the higher surface area to volume ratio in the polypropylene storage tube. This explanation would be consistent with observed PrP loss across multiple regimens of plastic exposure (see Figure 2). Notably, while aliquot size profoundly impacts PrP recovery from small (< 100 µL) aliquots, it does not appear to impact PrP levels in substantially larger CSF volumes. When comparing 1, 3 and 5 mL draws of a pooled CSF sample into identical 5 mL syringes, we did not see a difference in measured PrP (Figure S5B). The cylindrical shape of the syringe could also contribute to this finding, as the surface-area-to-volume ratio difference between different syringe volumes is less dramatic than that for very small sub-aliquots. These data have clinical implications: while downstream sub-aliquotting and storage can impact PrP levels, different syringe volumes during LPs performed with gentle aspiration will not greatly influence PrP recovery.

### Handling of test-retest samples.

We analyzed within-subject test-retest reliability of CSF PrP in four cohorts (Figure S7). Here is what we know about the handling history of these samples:

- Metformin trial placebo controls (Steven Arnold). Mean CV = 13% (Figure 3 and Figure S7A). *N*=18 samples comprise 2 lumbar punctures from each of 9 placebo-treated individuals from a randomized trial of metformin in individuals with mild cognitive impairment due to either Alzheimer disease or suspected non-amyloid pathology (SNAP). Test-retest interval ranged from 8 to 11 weeks. Lumbar punctures were performed fasting between 8:00a and 10:00a. CSF samples were handled according to a uniform protocol by the same staff, aliquotted into 0.5 mL aliquots within 1 hour of collection and then frozen on dry ice before storage at −80°C. The aliquots we received, approximately 1.75 years after the last sample was collected, were all 0.25 mL, indicating another round of freeze/thaw and aliquotting had occurred in the interim, but all samples were received in identical tubes with identical labeling.
- Sapropterin dihydrychloride trial participants (Kathryn Swoboda). Mean CV = 33% (Figure S7B). *N*=28 samples comprise 3 lumbar punctures from 8 individuals and 2 lumbar punctures from 2 individuals, all with Segawa syndrome (biallelic *GCH1* loss-of-function), enrolled in a trial monitoring effects of sapropterin dihydrochloride on CSF biomarkers. Test-retest interval ranged from 5 to 25 weeks. Lumbar punctures were performed at various times of day. Details of sample handling history are not known, but the aliquots we received were of various sizes (range: 150 µL to 1.3 mL) and were stored in different types of tubes (screw cap and flip top) with varied labeling (electronically generated and hand-written), suggesting a diverse sample handling history.
- MIND external lumbar drains (MGH MIND Tissue Bank). Mean CV = 40% (Figure S7C). *N*=18 samples comprise 3 days of external lumbar drains from 4 patients and 2 days of lumbar drains from 3 patients, with a test-retest interval ranging from 1 day to 4 months. These individuals were being evaluated at MGH for normal pressure hydrocephalus (*N*=7), *C. dificile* infection (*N*=1), or *Herpes simplex* infection (*N*=1). CSFs from these in-patient lumbar drains had contact with diverse plastics for varying amounts of time before freezing. In general, the samples passed through a pressure-measuring burette made of cellulose acetate propionate (CAP) before draining into a polyvinyl chloride (PVC) bag. CSF was later collected from the bag and frozen in either polystyrene (PS) or polypropylene (PP) tubes. Aliquots we received were of two different sizes: 0.5 mL and 4.0 mL.
- Pre-symptomatic and symptomatic PRNP mutation carriers (Michael Geschwind). Mean CV=34% in each (Figure S7D-E). Samples were collected between 2009 and 2017 at two sites (UCSF Parnassus NIH GCRC/CTSI and subsequently on the UCSF Mission Bay Neuroscience Clinical Research Unit) with multiple different physicians performing lumbar punctures according to a uniform protocol. Test-retest interval ranged from 2 months to 6 years. Samples were collected at various times of day and kept under refrigeration for variable amounts of time, ranging from a few hours to overnight, before being sent to UCSF CoreLabs. Samples collected prior to September 2016 were frozen immediately upon receipt at CoreLabs, and were later thawed and aliquotted in the first half of 2017. Beginning September 2016 CoreLabs aliquotted the samples upon receipt using polypropylene pipette tips (Rainin RT-L1000F) into 0.5 mL cryovials (Fisher 02-681-333) prior to first freeze. The sub-aliquots that we received were in identical tubes with uniform labels, and were all labeled as being 250 µL, however, we found that the actual recoverable volume in each tube varied, with some as low as 100 µL; all data reported here are from aliquots with at least 140 µL.

## Supplementary Table

**Table S1.** CSF samples analyzed. Abbreviations: normal pressure hydrocephalus (NPH); mild cognitive impairment with suspected nonamyloid pathology (MCI-SNAP).

| Cohort (Collaborator) | N | Diagnosis | Description |
| --- | --- | --- | --- |
| Metformin trial (Steven Arnold) | 18 | Alzheimer disease and MCI-SNAP | Placebo-treated controls from a randomized trial monitoring effects of metformin on CSF biomarkers<sup>32</sup>. 8-11 week test-retest. Samples were handled uniformly (see Supplementary Discussion) and were centrifuged prior to freezing. |
| MGH MIND Tissue Bank | 27 | NPH, *C. difficile*, herpes simplex | Large volume assay development samples from NPH patients (N=9), test-retest lumbar drains (N=18), and lumbar-thoracic gradient samples (N=8). Samples were centrifuged for 10 minutes at 2,000xG after receipt in our lab. |
| Sapropterin trial (Kathryn J Swoboda) | 28 | Segawa syndrome (*GCH1* loss of function) | Patients who received sapropterin dihydrochloride in a trial monitoring effects on CSF biomarkers (N=10 individuals). 5-25 week test-retest. Samples were centrifuged for 10 minutes at 2,000xG after receipt in our lab. |
| Bologna prion referrals (Piero Parchi) | 34 | Symptomatic prion and non-prion dementias | Dementia patients referred to the CJD Reference Center at University of Bologna due to suspected prion disease. Samples are autopsy-confirmed positive or negative for prion disease. Prion samples include sporadic and genetic (E200K, N=5). Prior to arriving at Dr. Parchi's lab from referring physicians, samples were variably centrifuged or not, and variably shipped frozen, cold, or at room temperature. Samples not marked as previously centrifuged were centrifuged for 10 minutes at 2,000xG after receipt in our lab. |
| Göttingen prion referrals (Inga Zerr) | 29 | Symptomatic prion and non-prion dementias | Dementia patients referred to the CJD Reference Center at University of Göttingen due to suspected prion disease. Samples are autopsy-confirmed positive or negative for prion disease. Prion samples include sporadic and genetic (D178N, N=2; E200K, N=2; V210I N=2). Samples were centrifuged for 10 minutes at 2,000xG after receipt in our lab. These samples were received after the data in Figure 1 were generated, so we added 0.03% CHAPS prior to sub-aliquotting and ELISA. |
| Cognitive impairment (Henrik Zetterberg) | 20 | Cognitive impairment | Patients with undiagnosed cognitive impairment and normal levels of CSF tau, phospho-tau, and amyloid beta. Samples were centrifuged for 10 minutes at 2,000xG after receipt in our lab. |
| UCSF (Michael Geschwind) | 61 | Symptomatic and pre-symptomatic genetic prion disease | Participants with *PRNP* mutations in the Early Diagnosis of Human Prion Disease study at UCSF<sup>55</sup>. The cohort includes N=61 samples from N=40 distinct individuals (28 pre-symptomatic and 12 symptomatic), with 1 to 5 samples per person collected at intervals ranging from 2 months to 6 years. Mutations represented include P102L (N=4 individuals), D178N (N=6), E200K (N=16), and ten other mutations (details omitted to protect patient privacy), including five with literature evidence for high penetrance and five without (see companion paper by Minikel et al). These samples were received after the data in Figure 1 were generated, so we added 0.03% CHAPS prior to sub-aliquotting and ELISA. Samples were never centrifuged. |
| **TOTAL** | **225** |  |  |

## Supplementary Figures

**Figure S1.**
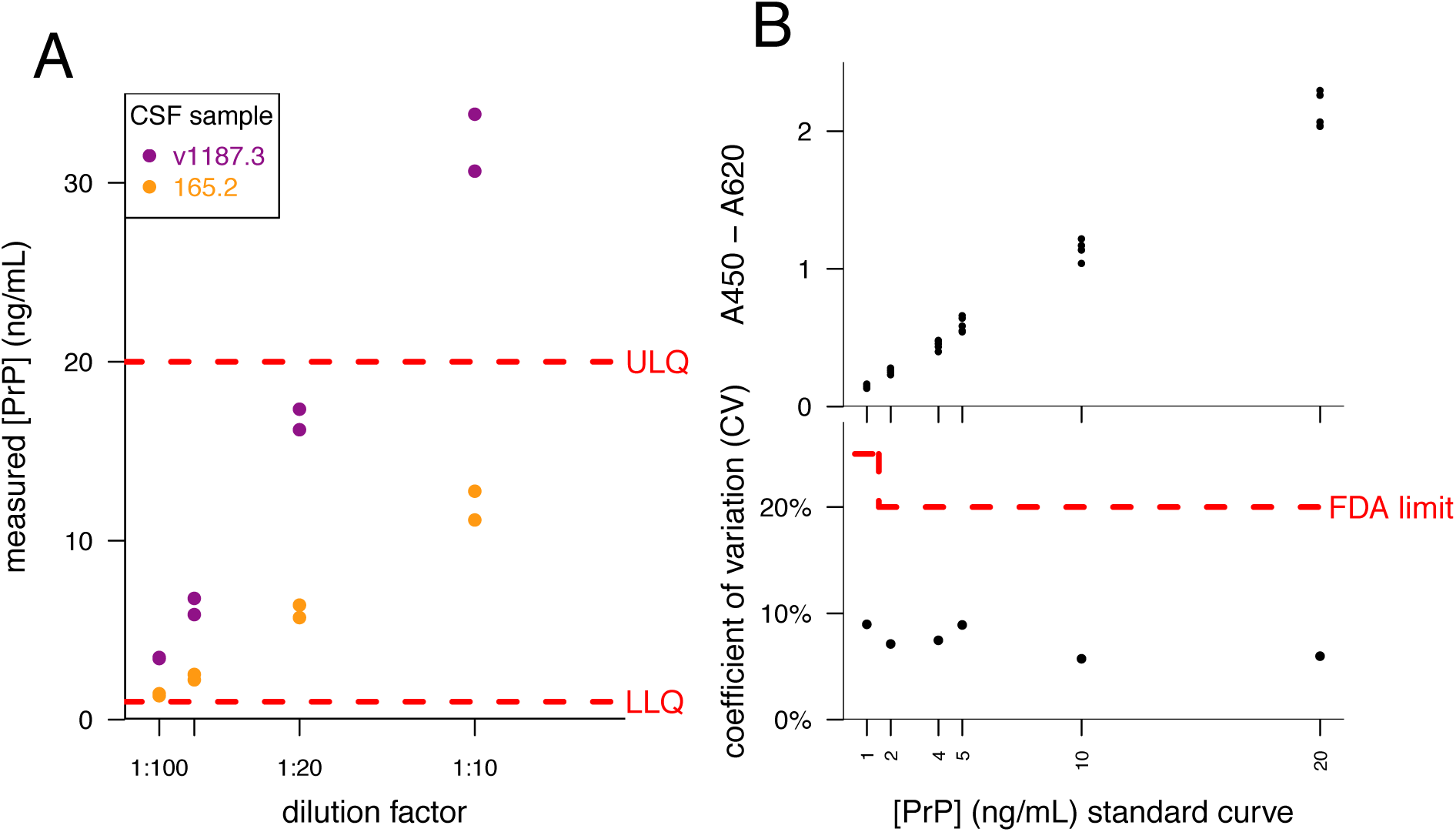
The BetaPrion^®^ Human PrP ELISA kit quantifies PrP in a technically reproducible and sensitive manner. A) Consistent dilution linearity was observed within the assay’s stated dynamic range of 1 – 20 ng/mL PrP, providing reassurance that this technique can be used to compare PrP levels across samples even when these levels differ by one log. Purple and yellow dots represent two different samples measured in duplicate at each of four dilutions. B) Five replicates of the kit’s internal six-point standard curve, reconstituted from lyophilized standards, were run in parallel on one plate. Across the dynamic range of the assay, the coefficient of variation falls below 10% for all points and well below the 20% FDA recommended limit in standard variability for ligand-binding assays.

**Figure S2.**
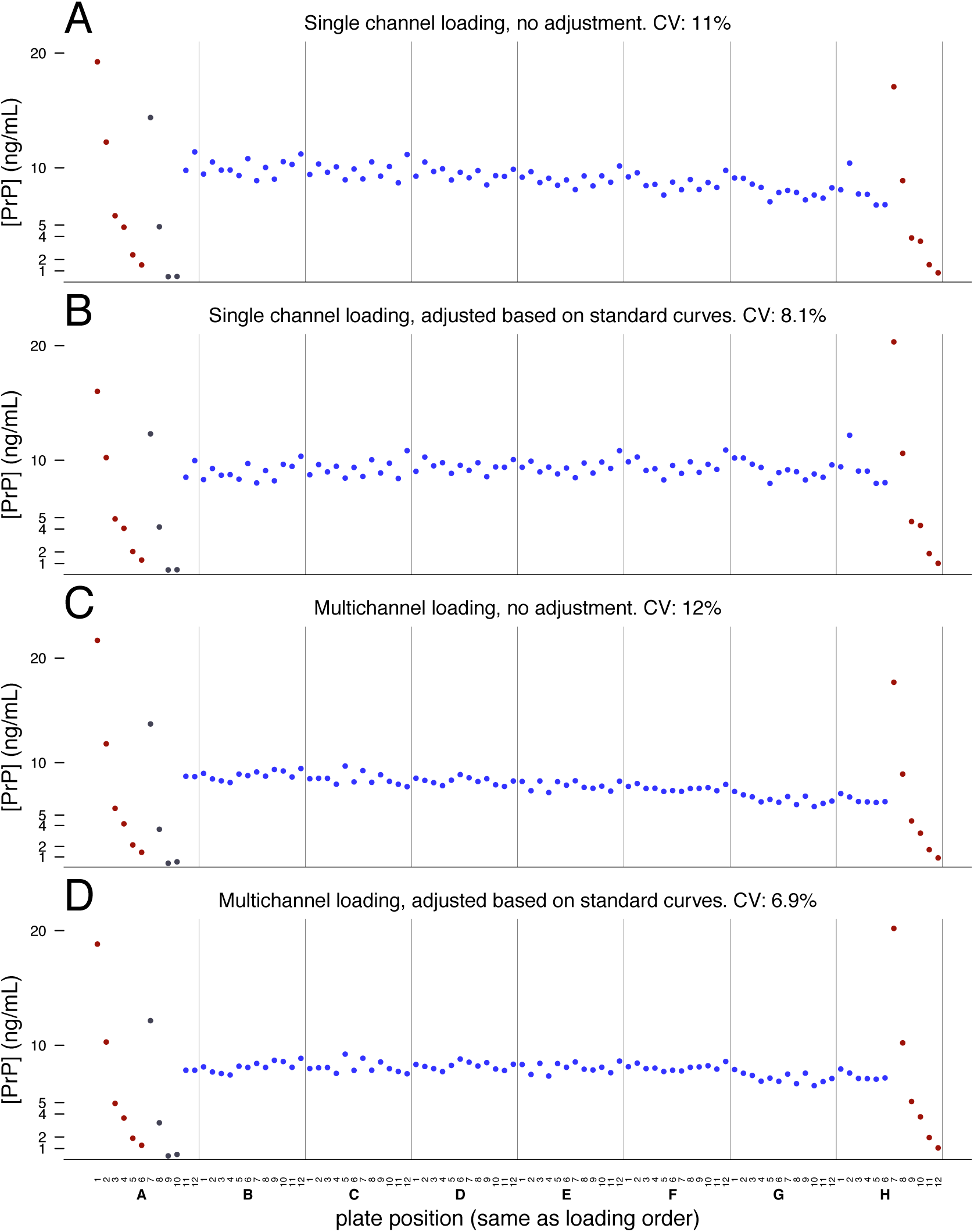
Plate position effects. Computed PrP levels for standard curves (red), kit controls (gray), or the CSF sample (blue) in two whole plates loaded with technical replicates of the same CSF sample (NPH sample v1209 with 0.03% CHAPS) using either a single channel pipette (A-B) or a multichannel pipette (C-D). Displayed are the unadjusted PrP values (A and C) or the PrP values after adjustment based on the difference between the standard curves at the beginning and end of the plate (B and D). See supplementary discussion for further interpretation.

**Figure S3.**
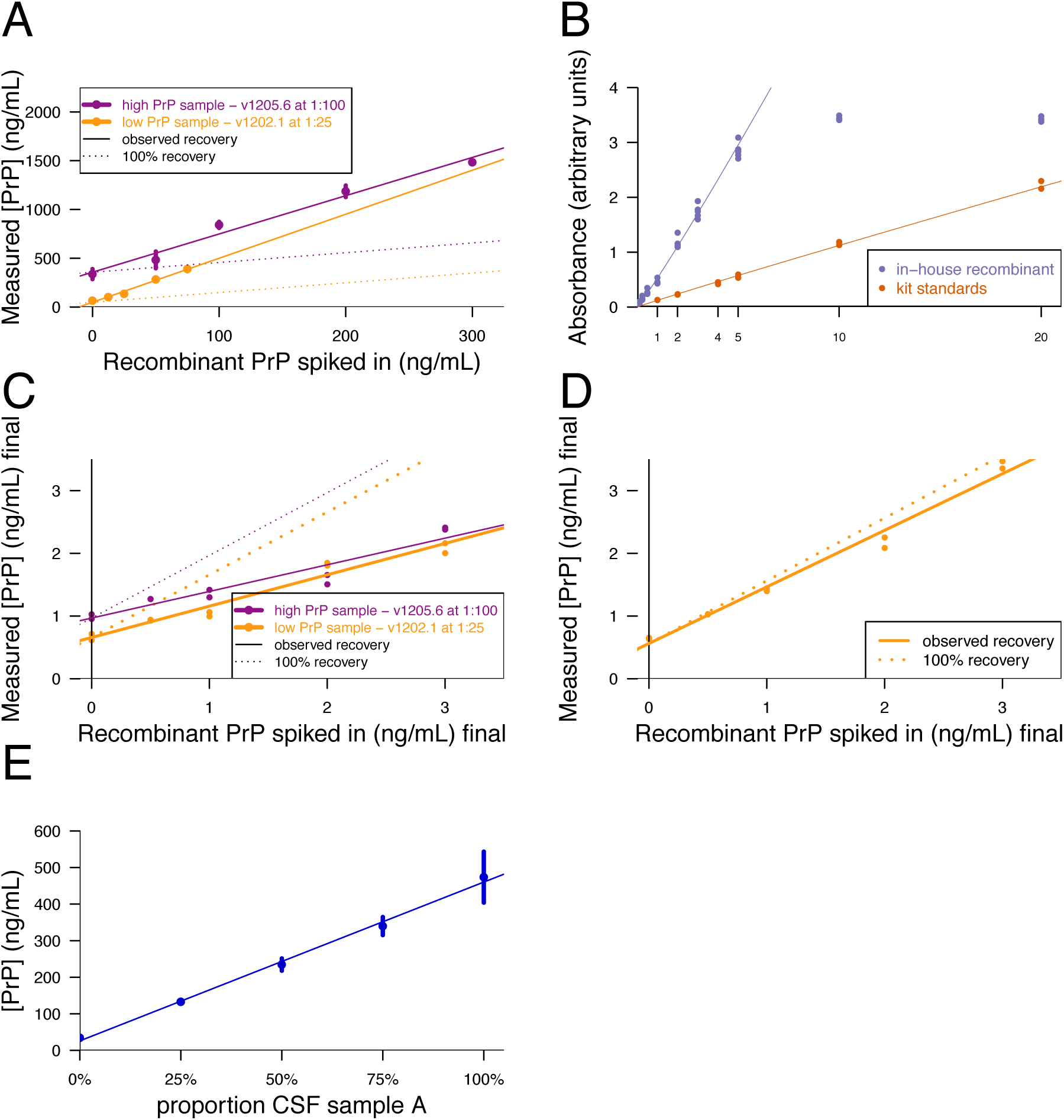
Spike recovery experiments. A) In-house produced full-length recombinant human prion protein, quantified by amino acid analysis (AAA) was spiked into two CSF samples previously established to have high and low baseline PrP. Recombinant PrP was over-recovered by 392-451% (meaning that measured concentrations were ~4x the expected concentrations) when compared to kit standards. B) A recombinant standard curve was prepared from AAA-quantified recombinant huPrP to match the nominal concentrations of each of the six points on the BetaPrion^®^ kit standard curve. Direct comparisons of the two series by ELISA showed the recombinant curve to be contain roughly 4x greater PrP at each point. C) Recombinant huPrP was measured according to a recombinant PrP standard curve. Recombinant PrP was diluted in CSF, then serially diluted into additional CSF to create a five-point series. The series of samples was re-frozen and measured by ELISA the next day. Under these conditions we observed 50.0% and 42.5% recovery for two different samples. D) The experiment in C was redone with the following modifications. Recombinant PrP was diluted directly in the initial aliquot tube with blocking buffer (5% BSA and 0.05% Tween-20 in PBS, filtered prior to use). It was further diluted in blocking buffer to prepare a series of solutions at 100x the desired final concentrations of points in the spike series. These samples were then added to CSF aliquots at a 1:100 concentration. These samples were then diluted in blocking buffer to their final plating concentration and measured in a same-day ELISA experiment. Under these conditions we observed 90.2% recovery. E) A high-PrP CSF sample (sample A) was titrated into a low-PrP CSF sample at varying ratios, with minimal CSF handling. We observed linear recovery of PrP. See supplementary discussion for further interpretation.

**Figure S4.**
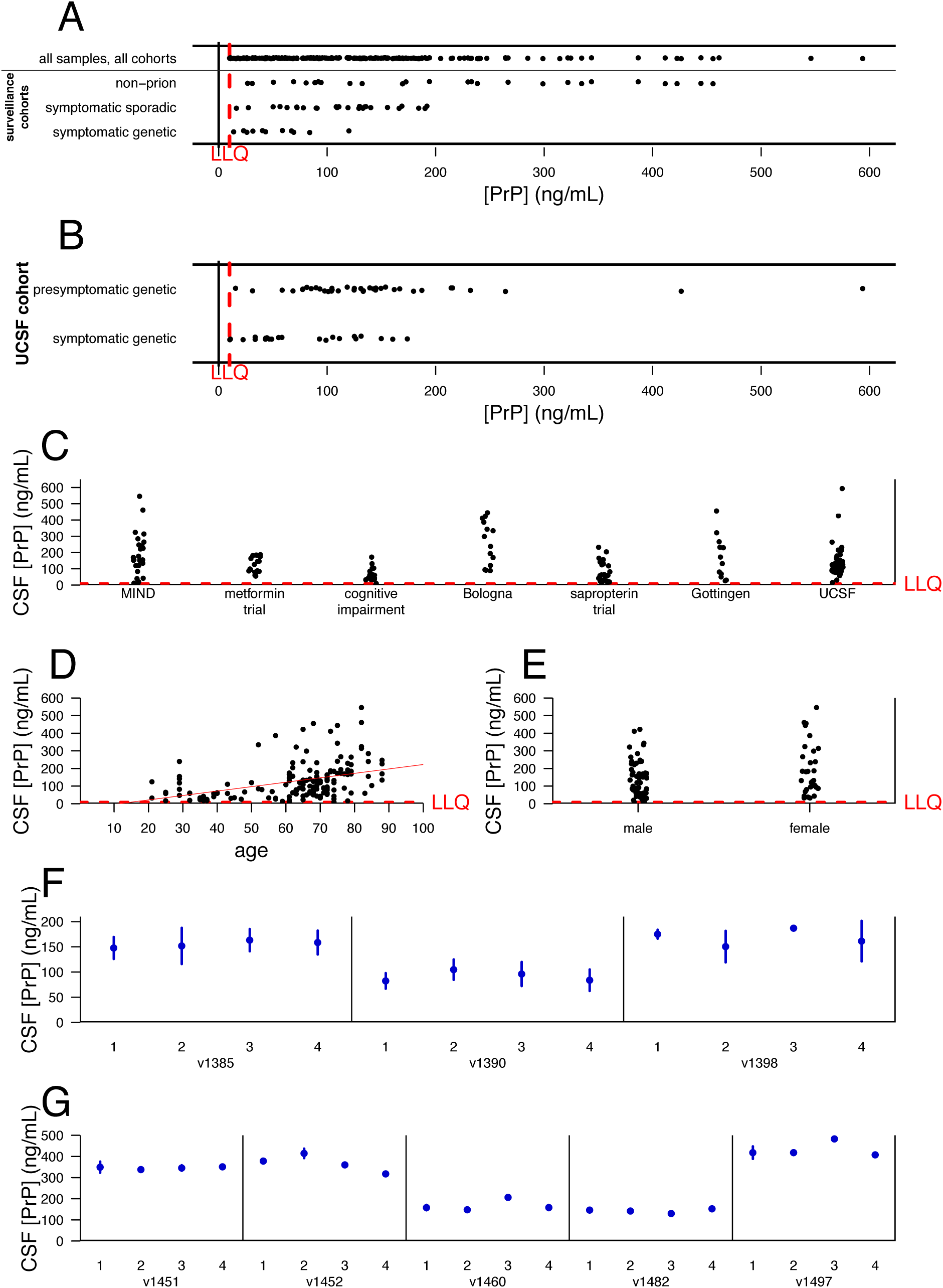
Candidate explanations for variability in CSF PrP levels. A) Within cohorts of individuals referred with a possible diagnosis of prion disease (Göttingen and Bologna cohorts), PrP levels are lower in individuals with prion disease than in individuals with other diagnoses. PrP levels in sporadic prion disease CSF average 42% of non-prion samples (P = 0.0001, Kolmogorov-Smirnov test) and in genetic prion disease CSF average 19% of non-prion samples (P = 2.6e-6, Kolmogorov-Smirnov test). B) Among individuals with a PRNP mutation (UCSF cohort), PrP levels in symptomatic individuals average 53% of those in pre-symptomatic individuals (P = .001, Kolmogorov-Smirnov test). C) CSF PrP levels vary dramatically between different cohorts in our study, even after excluding individuals with symptomatic prion disease (P = 1.1e-8, Type I ANOVA). D) CSF PrP is positively correlated with age (r = 0.47, P = 1.9e-9, Spearman rank test), although among our samples age is confounded with cohort, diagnosis, and likely with other unobserved variables, so it is unclear whether this correlation is biologically meaningful. For example, consider symptomatic prion disease patients in the two prion surveillance cohorts (Bologna and Göttingen). Symptomatic genetic patients were on average younger than symptomatic sporadic patients (mean 55 vs. 68 years old, P = 0.001, Kolmogorov-Smirnov test), and controlling for genetic vs. sporadic diagnosis eliminated any trend towards correlation between age and CSF PrP (linear regression, P = 0.37 with diagnosis as covariate, P = 0.04 without). E) Excluding individuals with symptomatic prion disease, CSF PrP does not differ between men and women (P = 0.31, Kolmogorov-Smirnov test). F) CSF PrP exhibits no lumbar-thoracic gradient within ~30 mL intrathecal CSF drips. From each of three individuals with normal pressure hydrocephalus, 29-32 mL of intrathecal CSF was collected via drip in 4 polystyrene tubes of 7-8 mL each, with “1” being the first tube and “4” being the final tube. Because CSF from further up the spinal column is expected to drain downward as CSF is removed, “1” represents the most lumbar CSF while “4” is the most thoracic. PrP exhibits no trend across tubes (P = 0.81, linear regression). Error bars show technical replicates performed in duplicate. G) CSF PrP likewise exhibits no lumbar-thoracic gradient when ~20 mL of CSF is drawn using gentle aspiration with a 24G Sprotte needle. Approximately 5 mL of CSF was drawn in each of four syringes; again, “1” is the most lumbar and “4” is the most thoracic. These samples included individuals diagnosed with Alzheimer’s disease, Parkinson’s disease, and undiagnosed individuals. PrP exhibits no trend across syringes (P = 0.93, linear regression). Error bars show technical replicates performed in duplicate.

**Figure S5.**
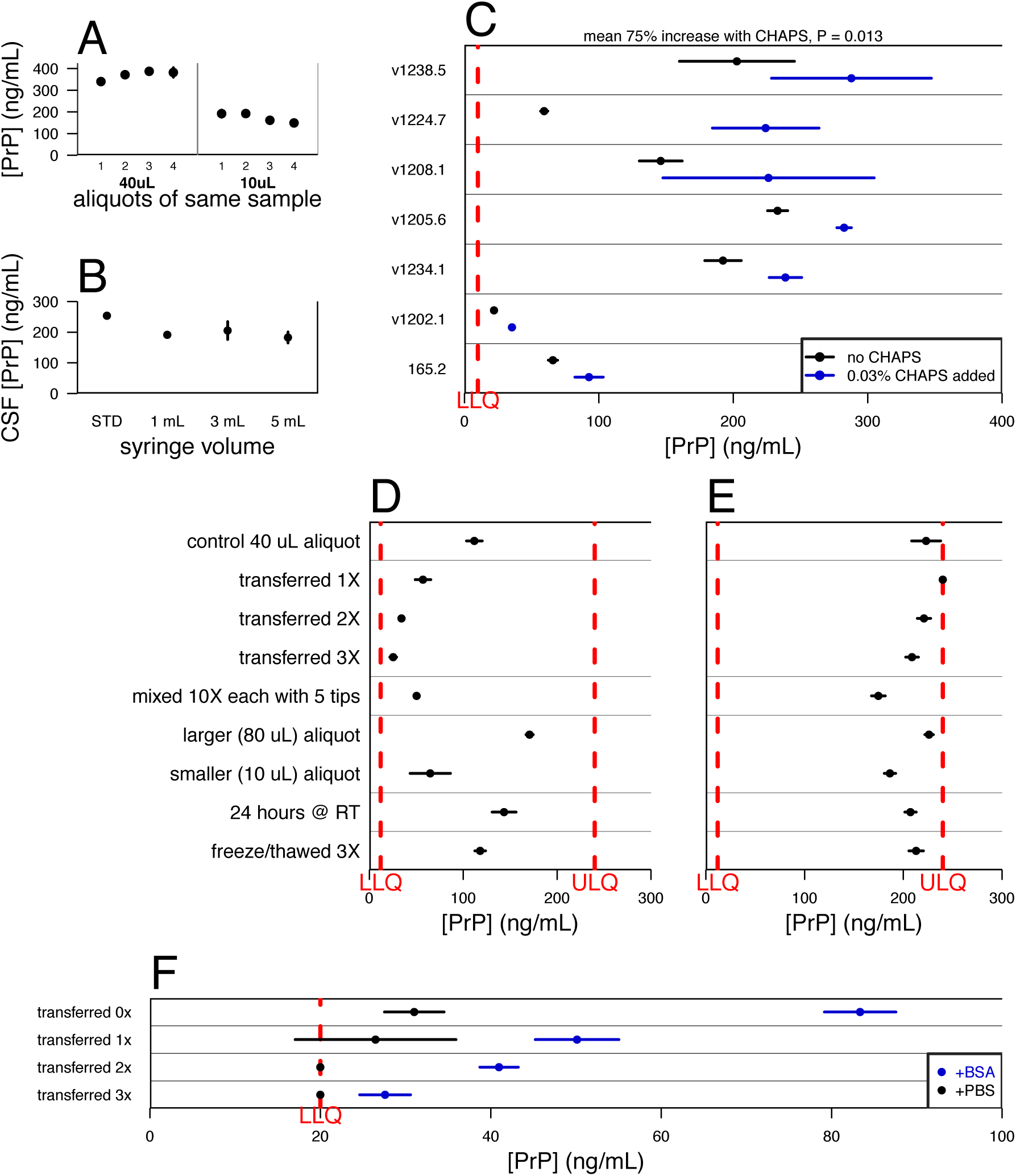
Additional evidence for loss of PrP to plastic adsorption. A) Differently sized aliquots of sample v1187 appear to have different PrP levels. Each dot is the mean, and line segment the 95% CI, of two technical replicates on the same plate. These samples did not contain CHAPS. B) A pooled CSF standard (STD) was warmed to 37°C and various volumes (1 mL, 3 mL, or 5 mL) were drawn into identical 5 mL syringes using a 24G Sprotte needle and allowed to sit for 15 minutes before ejection into tubes, centrifugation, and aliquotting. Samples were handled identically except for the volume drawn into the syringe. See supplementary discussion. C) After aliquotting and freeze/thaw, CSF samples were diluted into blocking buffer neat (black) or after addition of a final concentration of 0.03% CHAPS to the original storage tube (blue). Addition of CHAPS resulted in a 75% increase in apparent PrP level. See supplementary discussion. D and E) Replication of the findings from Figure 1A-B. The data in Figure 1 were generated using CSF samples from two different individuals; to rule out the possibility that some other inter-individual difference, rather than CHAPS, explained the difference in plastic loss, we repeated the experiment but with a single CSF sample divided into two halves which were then aliquotted without (D) or with (E) 0.03% CHAPS, subjected to the same battery of perturbations and plated at the same dilution. Because CHAPS increases overall PrP recovery, some replicates in (E) are at the upper limit of quantification; nevertheless, the results recapitulate Figure 1. F) 1 mg/mL (final concentration) BSA (blue), or PBS as a control (black), were added to CSF sample 165.2, which had an initial total protein level at the low end of the distribution of our samples (measured at 0.22 mg/mL with PBS), bringing it up to a total protein level at the high end of our samples (measured at 1.15 mg/mL after BSA spike-in). BSA or PBS were added after centrifugation but prior to aliquotting at 40 uL and re-freezing. 4 tubes of each sample were subsequently thawed and diluted into blocking buffer for analysis. Total recovery of PrP is increased in the BSA-spiked samples, analogous to panel B, although BSA is less effective at mitigating loss upon further transfer between tubes (compare to Figure 2A).

**Figure S6.**
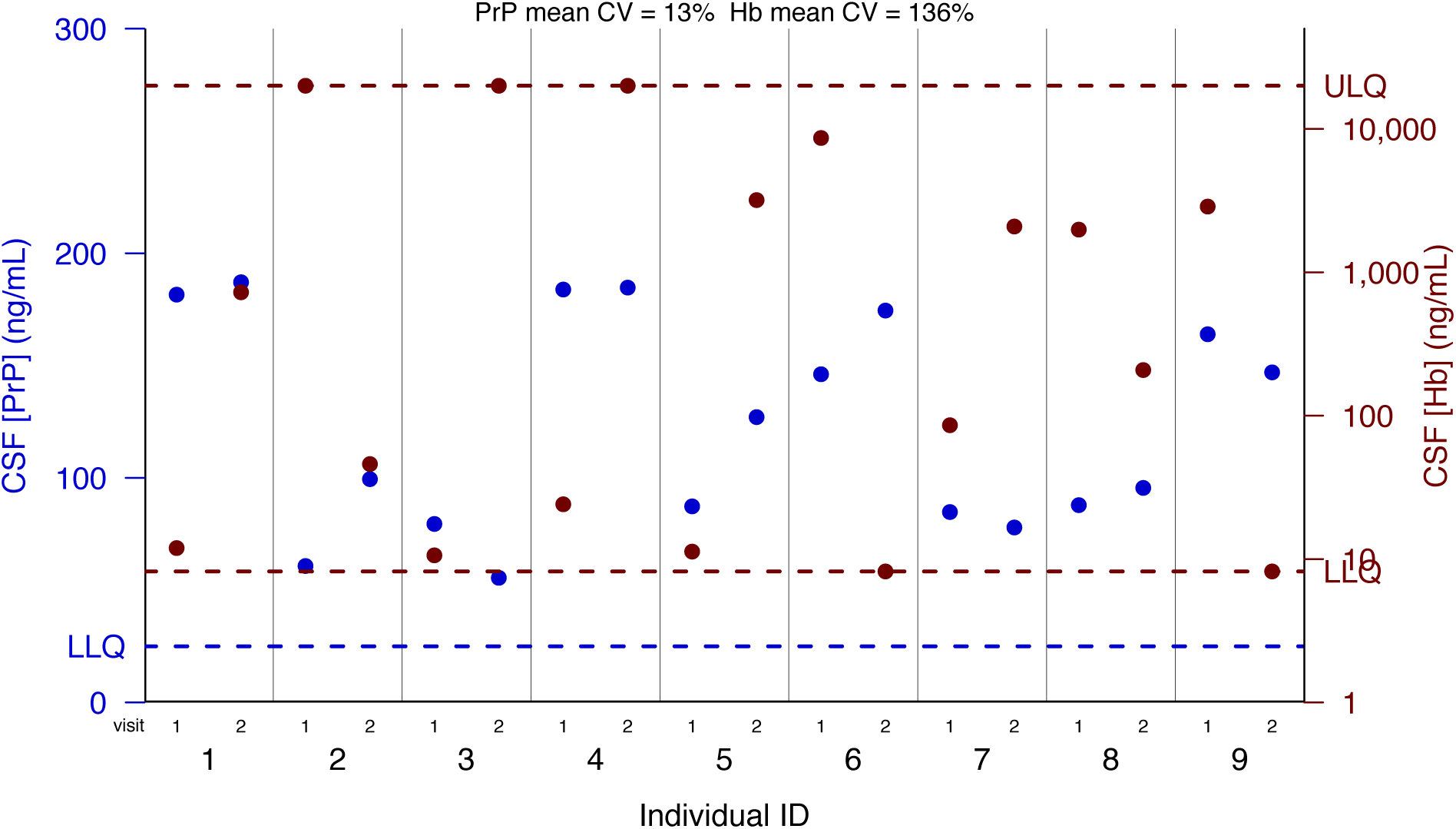
Hemoglobin in test-retest samples. Overlaid are PrP levels (blue, same data as shown in Figure 3) and hemoglobin levels (red) in test-retest samples. PrP exhibited good test-retest reliability (mean CV=13%) despite dramatic variation in hemoglobin (mean CV=136%), providing further evidence that blood contamination does not influence CSF PrP level.

**Figure S7.**
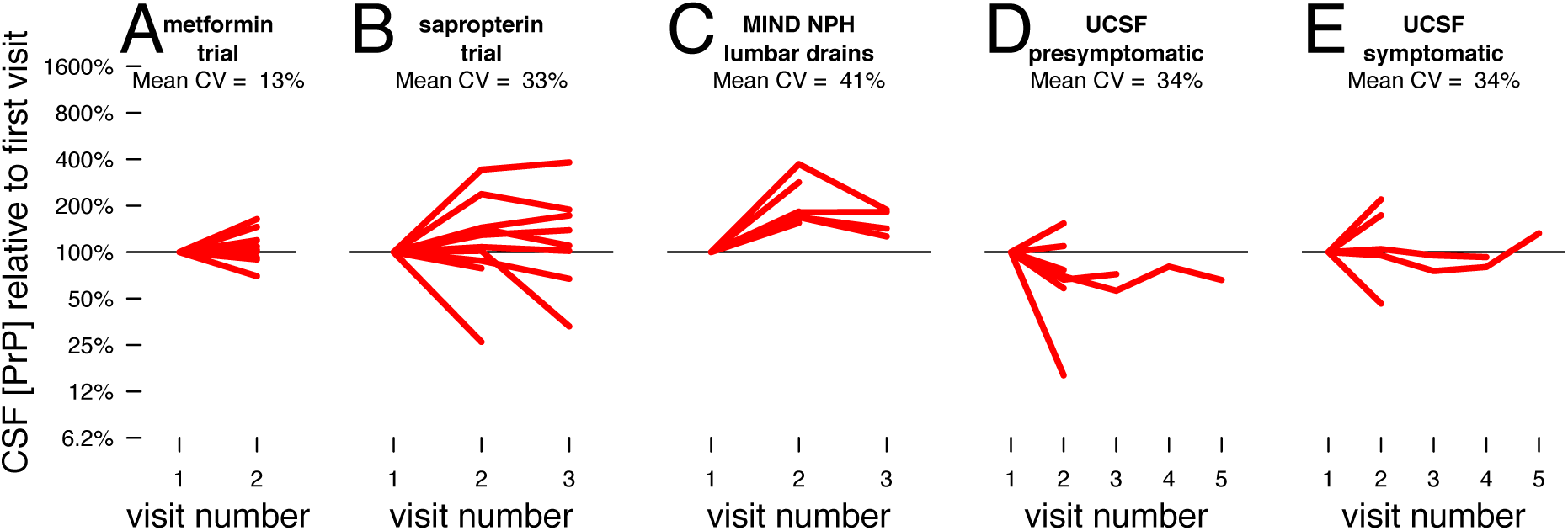
Test-retest reliability of CSF PrP in additional cohorts. Test-retest CSF PrP levels in A) metformin trial participants (Arnold) over 8-11 weeks, with mean CV=13% (same data from Figure 3 but plotted normalized to the PrP level at the first visit); B) sapropterin dihydrochloride trial participants (Swoboda) over 5-25 weeks, with mean CV=33%, C) NPH lumbar drains (MGH MIND Tissue Bank) over 1 day to 4 months, with mean CV=40%, D) pre-symptomatic and E) symptomatic PRNP mutation carriers (Geschwind) over 2 months to 6 years, each with mean CV=34%. The repeated 34% is not an error: the mean CVs in (D) and (E) happen to be the same (34.28% and 34.25%). See supplementary discussion for details on sample handling in these cohorts.

**Figure S8.**
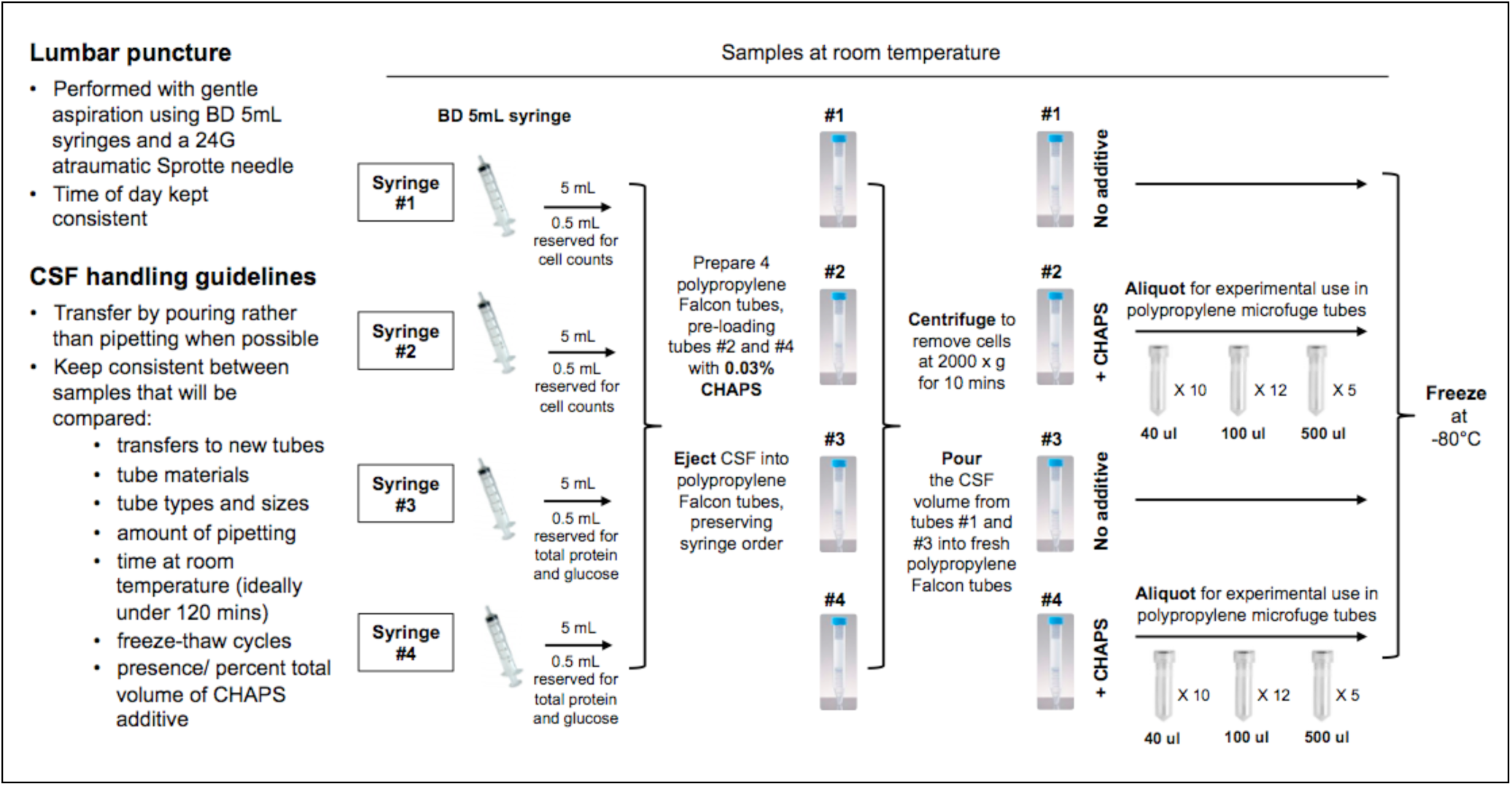
Protocol for collection of CSF for PrP measurement. We have incorporated our findings into the above protocol, which we are using to collect test-retest CSF for the purposes of PrP measurement in our ongoing clinical study.

